# Social context and dominance status contribute to sleep patterns and quality in groups of freely-moving mice

**DOI:** 10.1101/529669

**Authors:** Stoyo Karamihalev, Cornelia Flachskamm, Mayumi Kimura, Alon Chen

## Abstract

In many socially-living species, sleep patterns are subject to group influences, as individuals adjust to the presence, daily rhythms, and social pressures inherent to cohabitation. Disturbances in social functioning are comorbid with sleep problems in many prevalent psychiatric disorders, most notably, autism-spectrum, mood, and anxiety disorders (e.g., [1-3]). Our understanding of the common causality and the interplay between sleep impairment and psychiatric symptomatology could greatly benefit from experimental paradigms that allow simultaneous assessment of both domains of functioning. In laboratory mice, much is known about the sensitivity of sleep quality to a variety of experimental manipulations. Due to existing methodological restrictions, however, sleep studies are typically conducted in single-housed mice, thereby neglecting the influence of social dynamics and group-derived individual differences (however, see [4,5]). Here, we investigated sleep in a semi-naturalistic environment with freely-moving socially-housed groups of male mice using wireless electroencephalographic (EEG) monitoring devices and automated video tracking. In addition to multiple days of continuous behavioral data, we collected over fifty hours of EEG signal per mouse, recording simultaneously from all individuals in a group. We found evidence of in-group synchrony of sleep state patterns. Moreover, social status was a powerful predictor of sleep quality, such that sleep fragmentation, slow-wave sleep power, and rapid eye movement (REM) episode duration differed as a function of dominance status in the group. Finally, acute stress exposure had differential effects on REM sleep during recovery in dominant versus subordinate individuals. These findings highlight the importance of exploring sleep in a social context and are a step toward more informative research on the interplay between social functioning and sleep.

## Results

In order to study behavior in group-living mice, we used the “Social Box” (SB) paradigm, wherein a group of mice live together in an enriched environment under continuous video observation (Figure 1A; Movie 1) [6,7]. An automatic tracking system recorded the movement of each mouse in the SB, allowing us to detect and quantify the number, directionalities, and types of social interactions they displayed (*see Methods*). Wireless EEG and electromyographic (EMG) recordings were acquired from each mouse in the group simultaneously at several timepoints, including both dark phase and light phase recordings (the first 4 h of high-quality data collected in a 5 h recording period; Figure 1B-C, Supplementary Fig. 2).

**Figure 1.**
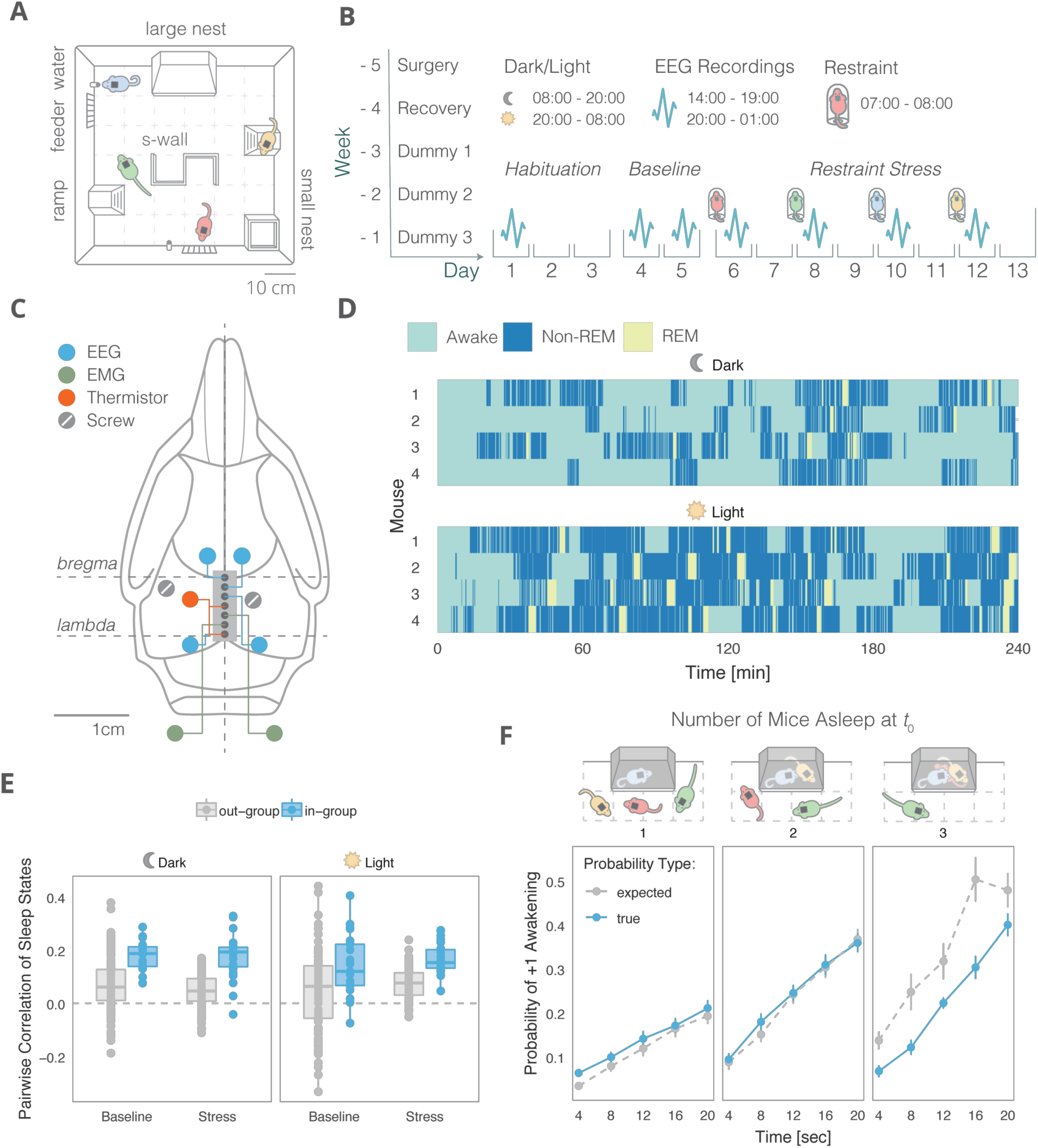
Sleep coordination in freely moving groups of mice. **(A)** The “Social Box” (SB) paradigm (described in detail in [6-8]). Each SB is an arena where a group of individuals cohabitate under continuous video monitoring for the duration of the experiment (several days). Each box contains a closed nest, a small open shelter, two ramps, two feeders, two water bottles, and an S-wall. **(B)** Experimental timeline. Implantation of EEG and EMG electrodes was performed five weeks before introduction to the SB. The surgery was followed by a week of recovery and three training weeks of habituating the animals to dummy head stages of increasing weights. Baseline behavior and EEGs were collected for 5 days, followed by individual stress (1 h restraint) before the beginning of the dark phase. A different animal from the same group was stressed every other day. **(C)** The coordinates of each recording electrode. Polygraphic signals were collected from four EEG channels, two EMG channels, and a thermistor. **(D)** Representative hypnograms from a single group of four mice (dark and light phase recordings, 14:00 – 19:00 & 20:00 – 01:00, resp.). **(E)** Pairwise correlations of sleep states. In-group sleep state correlations are higher than out-group correlations, indicating in-group sleep synchrony (in-group vs. out-group, *p* < 0.001). **(F)** Push-pull group effects on sleep. The probability of a mouse awakening within 5 epochs (20 seconds) if it is the only mouse in a group asleep at *t*_0_ is higher than expected from out-group measurements. Conversely, with three mice asleep at *t*_0_, the probability of one of them awakening in the same time frame is significantly lower than expected by chance (interaction of group type and number of mice, means +/- SEMs).

**Figure 2.**
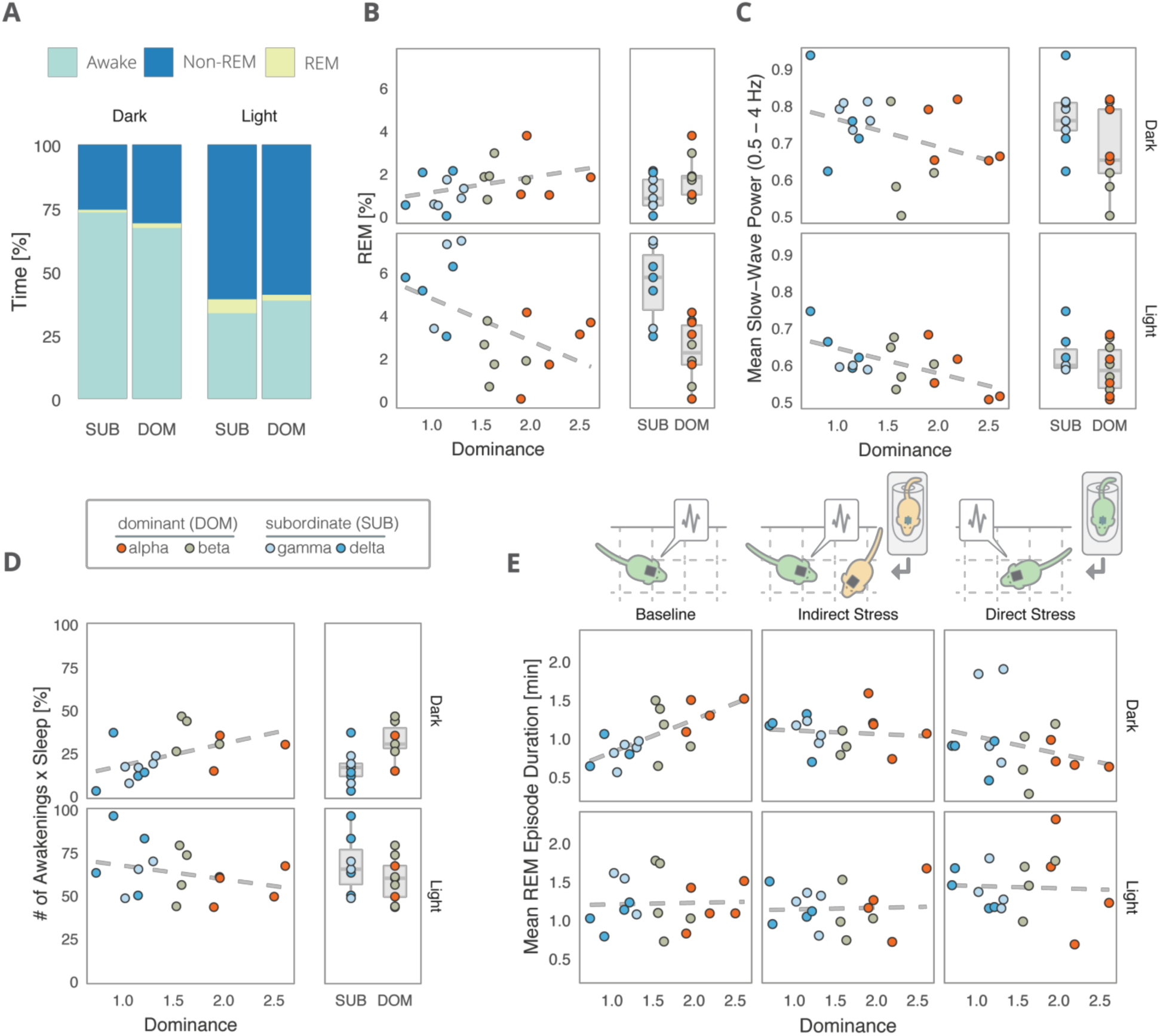
Social dominance status predicts sleep characteristics. **(A)** Average percent of time spent in each sleep stage differs between subordinate (SUB, ranks gamma and delta) and dominant (DOM, ranks apha and beta) animals, adjusted for between-group differences (approx. 4h of recording time per individual per light phase). **(B)** Baseline dominance (David’s Score rank) predicts increased group-adjusted dark phase REM sleep and decreased light phase REM sleep. **(C)** Lower-ranking individuals showed higher slow-wave activity during NREM sleep, suggesting baseline dominance levels predict group-adjusted mean NREM slow-wave power. **(D)** Sleep fragmentation, corrected for total sleep amount and group belonging, is increased in dominant animals compared to subordinates during the dark phase. **(E)** Social dominance mediates the effects of stress on group-adjusted mean REM episode duration.

### In-group correlation of sleep states

The ability to simultaneously collect sleep data from multiple group-housed individuals gave us the unique opportunity to investigate in-group temporal sleep synchrony (Figure 1D). In order to quantify sleep state synchrony, we dichotomized the scored sleep data into awake or asleep (either REM or NREM) and calculated the pairwise correlations in these sleep states between animals over time. To account for predictable circadian sleep patterns that occur innately in mice, independent of group belonging, we compared pairwise correlations for mice belonging to the same group (in-group) against out-of-group (out-group) correlations. During both the dark and light phase, baseline in-group correlations of sleep states were significantly higher than out-group correlations (*F*(1, 146) = 82.229, *p* < 0.001), suggesting inherent coherence in the group’s sleep dynamic (Figure 1E). Interestingly, the extent of these correlations does not appear to change during the stress phase of the experiment (following 1 h of acute restraint stress, Figure 1B) in a group type-specific manner (group type x stage interaction, *F*(1, 146) = 0.093, *p* > 0.05).

To further explore the baseline sleep cohesion in the group, we calculated the average probability of one or more additional animals waking up within 5 epochs (20 seconds) of an individual’s awakening at timepoint zero (t_0_, Figure 1F). We accounted for group influences by conditioning these probabilities on the number of individuals asleep at *t*_0_. We found that if only one mouse was asleep in a group at a given time, the true probability of it waking up within the allotted time interval is significantly higher than the expected probability (based on a comparison against out-group predictions). Conversely, if only one individual is awake at *t*_0_, the true probability of one of the three other individuals waking up is significantly reduced (probability type x N mice, *F*(2, 398) = 9.888, *p* < 0.001). When half of the group is asleep at *t*_0_, the true and expected probabilities of an additional awakening overlap. These results suggest that social context has a push-pull effect on an individual’s sleep pattern, wherein a mouse is more likely to wake up if its conspecifics are awake and more likely to stay asleep if its conspecifics are asleep.

### Dominance status contributes to sleep patterns

Social dominance status was assessed based on the numbers and directionality of aggressive interactions displayed during the first 12 h dark-phase recording. A dominance score (David’s Score, *see Methods*) was assigned to each mouse [6,8]. Additionally, ranking this score allowed for the separation of individuals in each group into dominant (DOM, ranks alpha and beta) and subordinate (SUB, ranks gamma and delta). This binary split was created to allow for equal sample sizes for each dominance status without segregating the individuals into too many groups, however we additionally look at correlations with the David’s Score directly. The assignment of dominance rankings based on the first night, rather than the entire monitoring period, made it possible for us to draw conclusions about the predictive power of social rank.

The average sleep state proportions for SUB and DOM individuals during both the light and dark phase recordings are summarized in Figure 2A. Curiously, while dominance contributed significantly to the proportion of REM sleep both in the dark and light phases, the association was positive in the dark phase and negative in the light phase (Figure 2B, rank x light phase interaction, *F*(1, 14) = 24.58, *p* < 0.001). We speculate that the decrease in REM sleep seen in the dark phase of SUB individuals could be interpreted as the acute suppression of REM following the stress associated with negative social interactions during the dark phase [9]. This may be followed by REM rebound during the light phase.

Slow-wave (0.5-4 Hz) power during NREM sleep reflects sleep intensity and homeostasis [10]. We found that the median slow-wave power during NREM sleep differs along dominance ranks, such that socially dominant individuals had overall reduced slow-wave activity (*F*(1, 13) = 5.045, *p* = 0.043), an effect which was especially pronounced during the light phase (Figure 2C, *F*(1, 11) = 6.37, *p* = 0.028). Concordant with this, we found sleep fragmentation to be higher in DOM individuals during the dark phase (Figure 2D, *F*(1, 14) = 6.04, *p* = 0.028 for post-hoc pairwise comparison prompted by a significant interaction effect of rank and light phase on sleep fragmentation, *F*(1, 12) = 7.57, *p* = 0.017).

In order to test if the relationship between social dominance and sleep properties is sensitive to stress, we compared sleep characteristics across social ranks at baseline with direct stress (1 h restraint) or indirect stress (another individual in the same group exposed to 1 h of restraint). A strong relationship between dominance and mean REM episode duration, present at baseline in the dark phase, was abolished if a stressed individual was present in the group, and reversed under direct stress (Figure 2E, Dominance x Mean REM episode duration in the dark phase, *F*(2, 26) = 5.73, *p* < 0.01). These findings suggest that the dominance effects on sleep may be disturbed by exposure to acute stress.

## Discussion

Social cohabitation is beneficial to mice and humans, offering, among others, protection from predation and opportunities for sharing of resources and parental care. These benefits, however, come at the expense of an individual having to adjust to group norms and compete with other group members.

Here we have shown that social cohabitation sets the sleep-wake behavior for group-living male mice. Given that a certain level of sleep synchronization seems to be the natural dynamic of a group, these findings hint at the possibility of psychiatric symptomatology manifesting as disturbances in synchronization to the sleep states of others. We hypothesize that, for example, mouse models of autism spectrum disorder would show impairment in their sensitivity to the sleep-wake patterns of their group-members. Likewise, while group sleep cohesion appears to increase under acute stress, we propose that chronic stress to an individual may disrupt both social relationships and sleep patterns sufficiently to uncouple them from the rest of their group.

Additionally, while cohabitation creates synchrony in overall sleep patterns, we have shown that group dynamics can also intensify individual differences. Social dominance is perhaps the most prominent and differentiating individual characteristic in male mice [8]. Here we show social rank-based differences in sleep architecture and REM episode duration, as well as slow-wave activity during NREM sleep. The pronounced increase in light phase REM sleep in combination with sustained slow-wave activity in subordinate mice may be indicative of exposure to stressful aggressive social interactions during the preceding active phase [9].

Simultaneous assessment of sleep and social behavior has uniquely allowed us to attempt a relatively detailed exploration of the connection between two separate, yet very much intertwined, domains of neurobiological functioning. We believe that this work emphasizes the importance of exploring sleep in a social environment and offers a way toward improved animal models of psychiatric disorders of sleep and social functioning.

## Movie 1

Movie 1. A representative clip from a Social Box recording. Four mice, implanted with EEG transmitters, move freely throughout the behavioral apparatus.

## Acknowledgments

Special thanks go to Yair Shemesh, Oren Forkosh, Noa Eren, Chadi Touma, and Markus Nussbaumer for their efforts in establishing the Social Box system. Thanks to Jessica Keverne for English writing support and advice. A.C. is the head of the Max Planck Society–Weizmann Institute of Science Laboratory for Experimental Neuropsychiatry and Behavioral Neurogenetics. This work is supported by: an FP7 Grant from the European Research Council (260463, A.C.); a research grant from the Israel Science Foundation (1565/15, A.C.); the ERANET Program, supported by the Chief Scientist Office of the Israeli Ministry of Health (3-11389, A.C.); the project was funded by the Federal Ministry of Education and Research under the funding code 01KU1501A (A.C.); research support from Roberto and Renata Ruhman (A.C.); research support from Bruno and Simone Licht; I-CORE Program of the Planning and Budgeting Committee and The Israel Science Foundation (grant no. 1916/12 to A.C.); the Nella and Leon Benoziyo Center for Neurological Diseases (A.C.); the Henry Chanoch Krenter Institute for Biomedical Imaging and Genomics (A.C.); the Perlman Family Foundation, founded by Louis L. and Anita M. Perlman (A.C.); the Adelis Foundation (A.C.); the Marc Besen and the Pratt Foundation (A.C.); and the Irving I. Moskowitz Foundation (A.C.).

## Author Contributions

Conceptualization, S.K. and A.C.; Methodology, C.F., M.K., and S.K.; Data analysis, S.K., and C.F.; Writing – Original Draft, S.K; Writing – Review & Editing, S.K., M.K., and A.C.; Funding Acquisition, A.C.; Supervision, A.C.

## Methods

### Animal housing and care

Male CD-1 (ICR) mice were bred and housed in an SPF-facility in temperature-controlled rooms under a 12h light/dark cycle with food and water available *ad libidum*. Upon weaning, mice were transferred into groups of four non-littermates and housed together until adulthood (>10 weeks of age). All animal studies were carried out in accordance with the European Community Council Directive. Animal experimental protocols were approved by the local commission for the Care and Use of Laboratory Animals of the Government of Upper Bavaria, Germany).

### Painting

All mice were marked prior to the start of an experiment to enable automatic video color tracking. Fur coloring was performed as described elsewhere [6,8]. Briefly, each mouse was painted in one of a set of four differently colored hair dyes (Tish & Snooky’s NYC Inc., New York) under mild isoflurane anesthesia using paint brushes. Excess dye was removed, and mice were single-housed for a recovery period of several hours after which they were re-introduced to their original groups.

### Wireless telemetry system

The wireless transmitters were custom-made by Multi Channel Systems GmbH (Reutlingen, Germany, Supplementary Fig. 1 A-B). Each transmitter weighed ca. 3 g and was attached to a seven-pin connector (Preci-Dip Durtal SA, Delémont, Switzerland). A detachable 100 mAh battery, weighing ca. 4 g, provided a maximum of five hours of continuous recording time at a sampling rate of 1 kHz (Supplementary Fig. 2). A receiver (Wireless 2100-RE, Multi Channel Systems GmbH, Reutlingen, Germany) was placed in a corner on top of the SB frame, such that any position in the social box was less that 1 m away. The receiver was connected to a computer via an interface board (Wireless 2100-IFB). Dedicated Multi Channel Systems software was used for acquisition (Multi Channel Experimenter). An electrode implant unit, consisting of a central seven-pin connector, was equipped with four soldered gold wire EEG electrodes, two gold wire EMG electrodes, and a thermistor (Tewa Temperature Sensors, Lublin, Poland), as described previously [11].

### Surgical procedures

Surgery was performed under inhalation anesthesia (mixture of isoflurane and oxygen) in a stereotaxic frame. Two EEG electrodes were inserted bilaterally anterior to bregma (+1.5 mm AP, ± 1 mm ML) and two more were inserted over the parietal cortex (AP −1 mm, ML ± 3 mm, Figure 1C). The thermistor was implanted unilaterally (AP −2 mm, ML 2 mm) for brain temperature monitoring. Additionally, two EMG electrodes were implanted into the trapezoid muscles. Finally, two anchor screws were placed into the skull for improved stability of the implant. The complete unit and screws were fixed to the cranial bone with dental resin. The entire surgical procedure lasted approximately 25 min per mouse. Prior to surgery, each animal received a subcutaneous injection of atropine sulfate (0.05 mg/kg, Atropine, Braun Melsungen, Melsungen, Germany) for stabilization of cardiovascular function and meloxicam (1 mg/kg, Metacam, Braun Melsungen, Melsungen, Germany) for analgesia. Meloxicam was additionally administered at 24 h and 48 h after surgery.

The surgery was followed by a week of recovery, during which time animals were housed in their original groups and bodyweights were monitored daily. For three weeks after the recovery week, the animals were acclimated to carry progressively heavier dummy-transmitter devices (custom-made using aluminum plates in the shape of the transmitter, dummy 1 – 2 g, dummy 2 – ca. 4g, and dummy 3 – ca. 7 g). Each dummy was worn continuously for a week in the three weeks leading up to the first measurement (Figure 1B, Movie 1). During the entire three-week training period, the animals were habituated to gentle daily handling.

### Data acquisition

At the start of each recording, each mouse was removed from the SB for several minutes. During this time, the dummy transmitter was replaced with a real transmitter with a charged battery attached. The mouse was then reintroduced into the SB. The transmitter and battery were replaced with the 4 g dummy the following day, which was worn until the next recording.

### Telemetric data processing and sleep-wake classification

Telemetric data was processed offline for analysis with a LabVIEW-based acquisition program (National Instruments, Austin, TX, USA), customized for use in mice (EGErAVigilanz, SEA Cologne, Germany) [13]. EEG and EMG signals were amplified 10000 times and filtered (EEG: 0.1-460 Hz, EMG: >200 Hz).

Sleep/wake states were determined manually by an experienced scorer simultaneously taking into account the parietal EEG as well as the EMG signal. Vigilance states were characterized as “awake”, “REM”, and “non-REM” (non-rapid eye movement sleep).

A Fast Fourier Transform (FFT) algorithm was used on data binned into four-second epochs for power spectral analysis. Mean values of the EEG spectrum/0.25 Hz were calculated and normalized per animal per recording using the individual mean of the total EEG power from all vigilance states across all frequency bins and epochs. Slow-wave activity during NREM sleep was assessed by summing over the power densities between 0.5 Hz and 4 Hz. This power was then averaged per animal across the measurement period [13].

### The Social Box and automated behavioral tracking

The SB is an enriched housing environment designed to house groups of mice living together over extended periods of time [6-8]. Each apparatus consisted of a 60 cm x 60 cm enclosure under a stationary video camera. LED lights were set on a 12 h schedule switching between light phase illumination (ca. 200 lux) and dark phase illumination (2-5 lux). Each SB contained a nest, two ramps, a S-wall, a small shelter, two feeders, and two water bottles (Fig. 1A). The behavior of mice in the SB were recorded continuously and tracked automatically using a specialized software written in Matlab (Mathworks Inc.).

### Assignment of dominance ranks

Agonistic interactions between mice throughout the SB monitoring period were captured and classified based on movement trajectories as described in detail elsewhere [6,8]. The David’s score was used as a continuous in-group measure of dominance, calculated based on the numbers and directionalities of aggressive chases [8,12]. In order to create discrete groups, mice were additionally ranked based on this score and the two top-ranking individuals were considered to be “DOM” while the 3^rd^ and 4^th^ ranking individuals were considered “SUB”.

### Statistical analyses

All statistical analyses were performed in R (www.R-project.org) [14]. Mixed effects modelling was aided by the “nlme” package [15]. All tests of social dominance against sleep properties included group belonging as a covariate factor in the analysis, thus adjusting for the effects of group differences. Accordingly, the values plotted for such comparisons are always adjusted for group effects.

## Supplemental Information

### Supplementary Figure 1

**Supplementary Figure 1.**
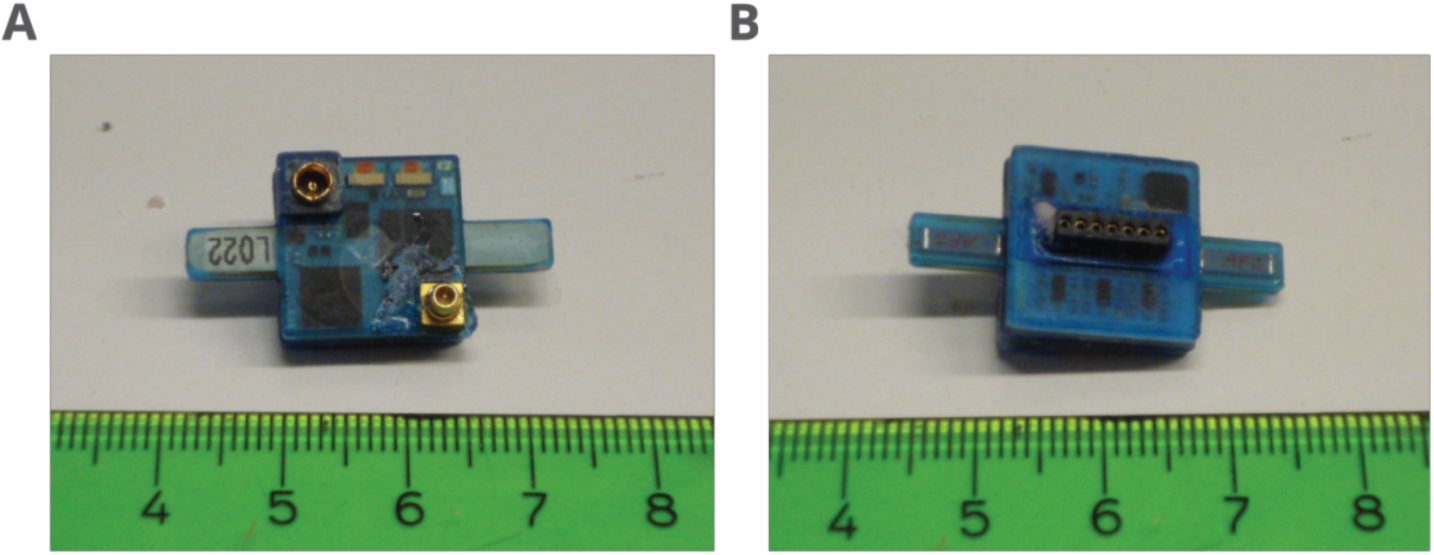
Related to Figure 1. View of the bottom **(A)** and top **(B)** of the custom-made wireless transmitter by Multi Channel Systems GmbH (Reutlingen, Germany).

### Supplementary Figure 2

**Supplementary Figure 2.**
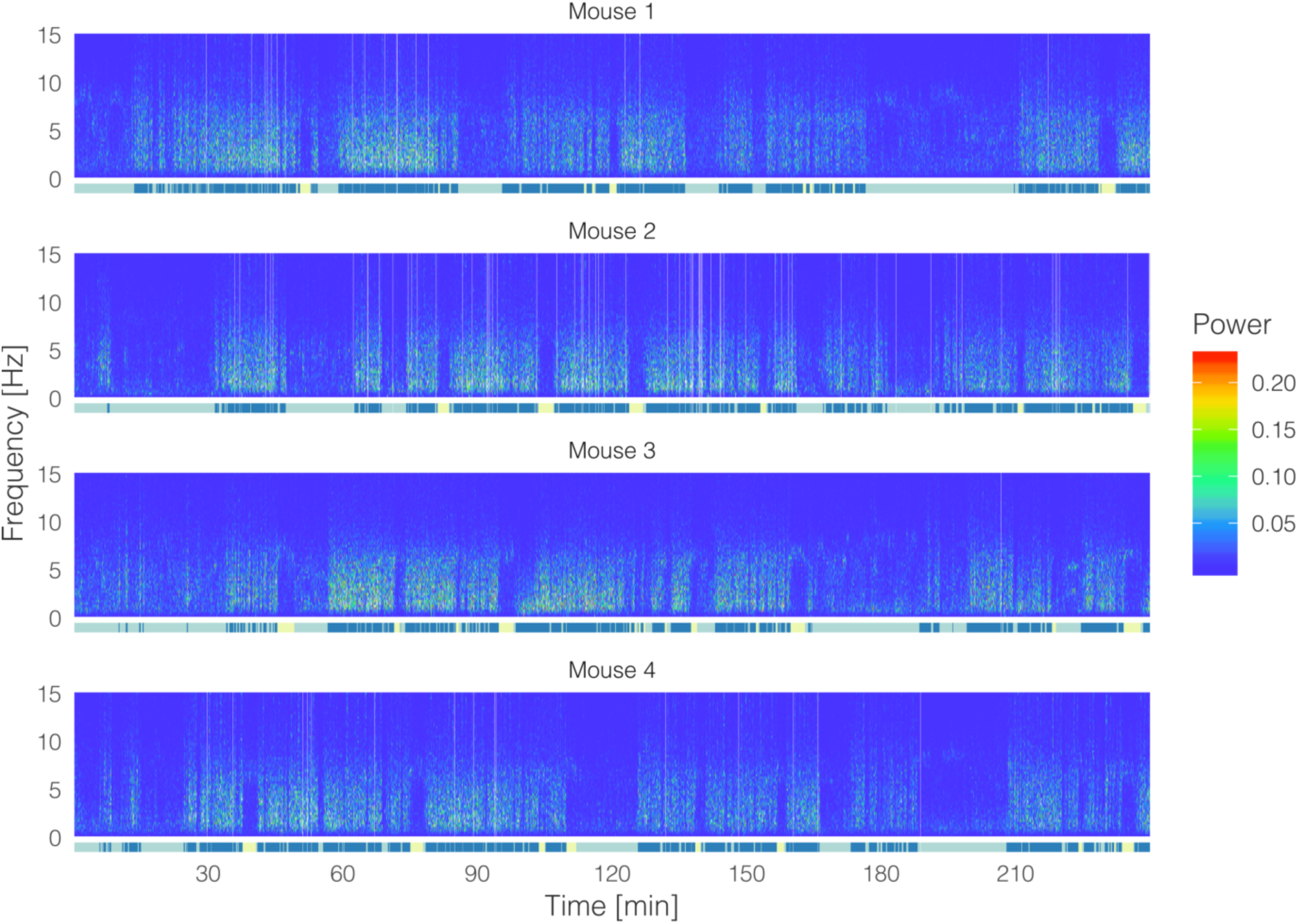
Related to Figure 1. Example spectrograms (with each hypnogram at the bottom) from a four-hour light-phase EEG recording in four freely-behaving group-housed mice acquired simultaneously in the Social Box. Color coded plots display differences in the intensity of EEG power densities.

